# Inhibition of KLK8 promotes pulmonary endothelial repair by restoring VE-cadherin/Akt/FOXM1 pathway

**DOI:** 10.1101/2023.12.06.570377

**Authors:** Ying Zhao, Hui Ji, Feng Han, Qing-Feng Xu, Hui Zhang, Di Liu, Juan Wei, Dan-Hong Xu, Lai Jiang, Jian-Kui Du, Ping-Bo Xu, Yu-Jian Liu, Xiao-Yan Zhu

## Abstract

The tissue kallikrein-related peptidases (KLKs) are secreted serine proteases deeply involved in angiogenesis. However, whether KLKs are involved in the regulation of endothelial regeneration during sepsis remains unknown. By comparing the mRNA levels of 15 KLKs, we found that KLK8 was the highest induced KLK member in lung tissues or primary isolated mouse lung vascular endothelial cells (MLVECs) exposed to lipopolysaccharide (LPS). Adenovirus-mediated overexpression of KLK8 caused endothelial hyperpermeability both *in vitro* and *in vivo*. Inhibition of KLK8, by either gene knockout or KLK8 neutralizing antibodies, alleviated sepsis-induced endothelial hyperpermeability, acute lung injury and mortality. Mechanistically, transcription profiling of KLK8-overexpressed endothelial cells revealed a central role of forkhead box M1 (FOXM1) downregulation in mediating the pro-injury and anti-proliferation effects of KLK8. KLK8 cleaved VE-cadherin and consequently suppressed FOXM1 expression by inactivation of the VE-cadherin/Akt pathway. KLK8 deficiency or blockade rescued VE-cadherin/Akt/FOXM1 pathway, thus promoting endothelium regeneration. This study reveals a critical role for KLK8-induced inactivation of VE-cadherin/Akt/FOXM1 pathway in mediating the impairment of endothelial regeneration and the consequent lung vascular leakiness in response to sepsis.

**Highlights:**

- Upregulated KLK8 mediates lung endothelial barrier dysfunction during sepsis
- KLK8 inactivates VE-cadherin/Akt/FOXM1, thus impairing endothelium regeneration
- KLK8 deficiency or blockade rescues VE-cadherin/Akt/FOXM1 signaling pathway
- KLK8 deficiency or blockade promotes endothelium regeneration during sepsis
- KLK8 deficiency or blockade attenuates sepsis-induced acute lung injury and mortality

## 1. Introduction

Sepsis, a devastating complication commonly observed in patients with serious infection, burns, or trauma, is associated with progressive multiple organ dysfunction syndrome (MODS), which significantly contributes to mortality (Font et al., 2020, Barichello et al., 2022). The microvasculature, particularly the endothelial cells lining the microvessels, is highly susceptible to the detrimental effects of sepsis (Joffre et al., 2020, Dolmatova et al., 2021). Endothelial dysfunction, a significant concern in sepsis, is encountered during various medical procedures such as fluid resuscitation and thrombolytic therapy, further exacerbating the condition (De Backer et al., 2021, Bakker et al., 2022, Molema and Zijlstra, 2022).

Among the organs affected by sepsis, the lung is particularly vulnerable (Genga and Russell, 2017, Meyer et al., 2021). Endothelial cells, acting as the first line of defense in sepsis pathogenesis, undergo intercellular connection disruption, morphological changes, and apoptosis, leading to pulmonary endothelial hyperpermeability and edema (Joffre et al., 2020, Vassiliou et al., 2020). Additionally, the presence of numerous toxins and inflammatory mediators in the circulation following sepsis directly impairs the proliferation ability of pulmonary microvascular endothelial cells, hindering the self-repair of microvessels and exacerbating sepsis-induced lung injury (Evans et al., 2021, Villar et al., 2019). Consequently, targeting endothelial repair mechanisms represents a promising treatment strategy to restore the integrity of the pulmonary microvascular barrier following sepsis.

The tissue kallikrein-related peptidases (KLKs) constitute a family of 15 homologous secreted serine peptidases encoded by a gene cluster located at chromosome 19q13.4 (Stefanini et al., 2015, Lenga Ma Bonda et al., 2018). Growing evidence suggests that KLKs play a significant role in angiogenesis by regulating endothelial cell tube formation, proliferation, and migration (Srinivasan and Kryza, 2022, Yao et al., 2013, Kryza et al., 2018). Moreover, certain KLKs, including KLK1, KLK5, KLK6, and KLK8, have been implicated in promoting endothelial-to-mesenchymal transition (EndMT) in vascular diseases (Yao et al., 2015, Du et al., 2021). However, the involvement of KLKs in the regulation of pulmonary endothelial regeneration and vascular repair during sepsis remains unknown.

In this study, we investigated the mRNA expression levels of all 15 mouse KLKs in lung tissues obtained from lipopolysaccharide (LPS)-treated mice, as well as LPS-treated primary isolated mouse lung vascular endothelial cells (MLVECs). Among the KLK family members, KLK8 exhibited the highest upregulation in lung tissues and MLVECs exposed to LPS. To elucidate the role of KLK8, we employed KLK8 knockout, intra-pulmonary KLK8-overexpressed, and KLK8 neutralizing antibodies-treated mice. The collective evidence demonstrated that upregulation of KLK8 contributed to pulmonary endothelial hyperpermeability and acute lung injury induced by LPS or cecal ligation and puncture (CLP)-induced sepsis. Transcriptional profiling revealed that the KLK8-induced inactivation of the vascular endothelial (VE)-cadherin/Akt/forkhead box M1 (FOXM1) pathway played a crucial role in mediating the impairment of endothelial regeneration and subsequent lung vascular leakiness in response to LPS or CLP-induced sepsis. These findings provide a potential target for the development of vascular protective drugs for sepsis.

## 2. Results

### 2.1 KLK8 is upregulated in vascular endothelial cells in lung tissues of endotoxemic mice or CLP-induced septic mice

To investigate the potential involvement of KLKs in the development of pulmonary endothelial barrier dysfunction, we initially examined the gene expression levels of KLK family members in lung tissues of endotoxemic mice and LPS-treated MLVECs. Our findings revealed a significant increase in pulmonary mRNA expressions of KLK2 and KLK8, while KLK1, KLK4, KLK5, KLK7, and KLK10 were decreased in the endotoxemic mice compared to the control mice (Figure 1A). Furthermore, LPS treatment at varying concentrations (10 ∼ 1000 ng/ml) resulted in dose-dependent increases in the mRNA levels of KLK2 and KLK8, along with decreases in KLK5, KLK7, and KLK10 in primary cultured MLVECs (Figure 1B). Notably, KLK8 exhibited the highest induction among the KLK family members in both lung tissues of endotoxemic mice (3.57-fold change compared to control) and LPS-treated MLVECs (3.44-fold change compared to control in 1000 ng/mL LPS-treated MLVECs).

**Figure 1.**
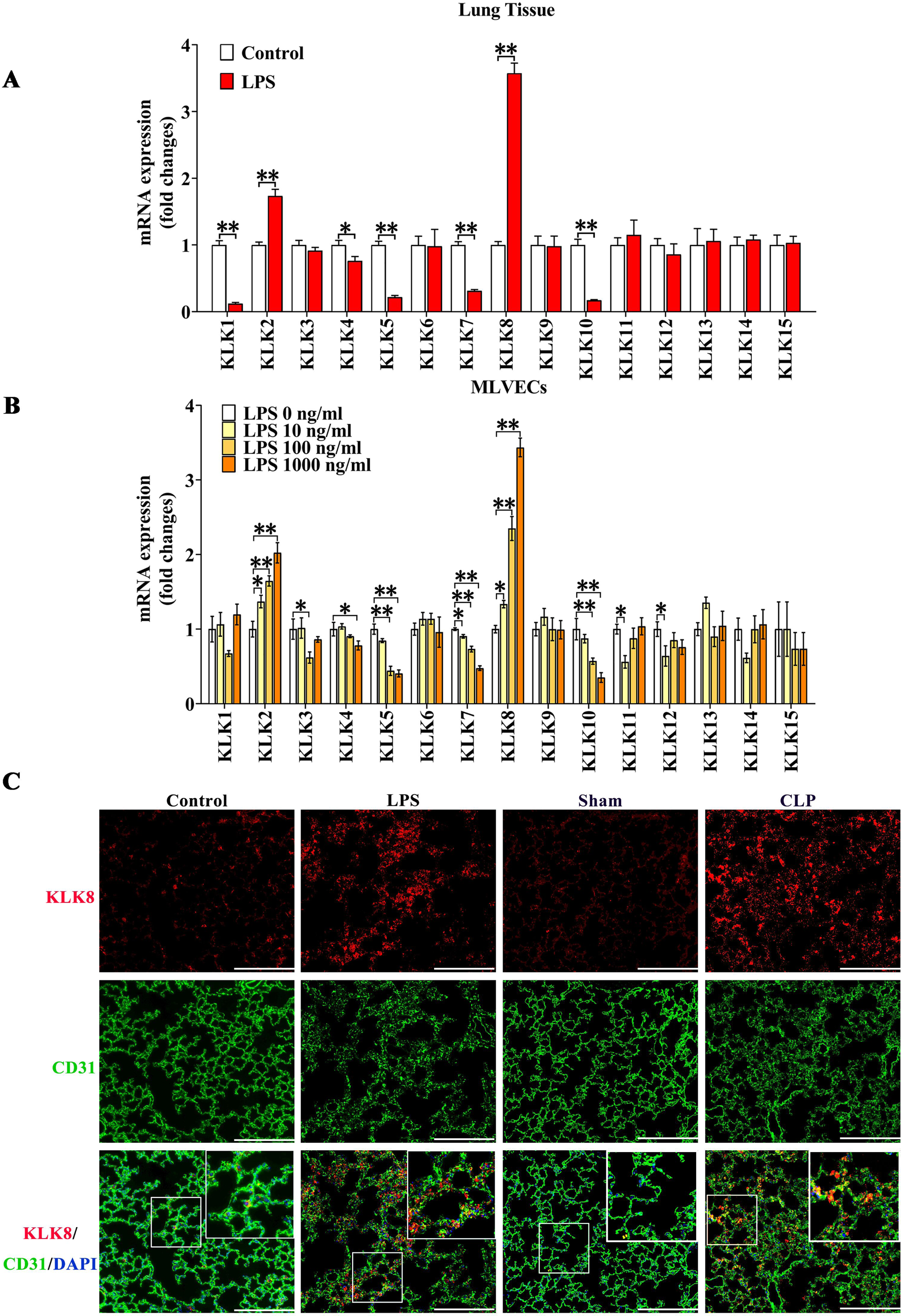
KLK8 is upregulated in vascular endothelial cells in lung tissues of endotoxemic mice or CLP-induced septic mice. **A-B,** Mice were intraperitoneally injected with LPS (30 mg/kg) or saline. MLVECs were treated with LPS at the indicated doses. The mRNA levels of KLK family members in the lung tissues (**A**, n=6) or MLVECs (**B**, n=4) exposed to LPS for 24 hours. **C**, Lung tissues obtained at 48 hours after LPS injection or CLP/sham operation were stained with fluorophore-labeled antibodies against endothelial cell marker CD31 (green) and KLK8 (red). Nuclei were counterstained with DAPI (blue). Scale bars correspond to 100 μm. Statistical analyses were performed using an unpaired, two-tailed Student’s t-test (A) or one-way ANOVA with Student-Newman-Keuls post hoc test (B). Data are presented as the mean ± SEM * p<0.05, ** p<0.01.

To determine the expression of KLK8 in pulmonary endothelial cells, we conducted double immunofluorescence staining using antibodies against KLK8 and the vascular endothelial cell marker CD31. Compared to the control and Sham mice, we observed a significant increase in KLK8^+^/CD31^+^ cells in lung tissue sections from both endotoxemic mice and CLP-induced septic mice (Figure 1C and supplemental Figure S1), respectively.

### 2.2 KLK8 overexpression leads to endothelial hyperpermeability and acute lung injury

We proceeded to investigate the impact of KLK8 on primary cultured MLVECs. Treatment of MLVECs with KLK8 adenovirus (Ad-KLK8) led to a dose-dependent increase in KLK8 expression (Figure 2A). The maintenance of endothelial barrier integrity relies on the tight-junction protein ZO-1 and the endothelial cell-specific adherens junction molecule CD31 (Zhuang et al., 2016). Additionally, ICAM-1, abundantly present on activated endothelial linings, serves as a marker for inflammation-associated endothelial dysfunction (Bui et al., 2020). As shown in Figure 2A&B, Ad-KLK8 treatment dose-dependently induced endothelial cell damage, as evidenced by decreased CD31/ZO-1 protein levels, reduced cell viability, and increased ICAM-1 expression. Furthermore, Ad-KLK8 treatment significantly increased the permeability of MLVECs (Figure 2C).

**Figure 2.**
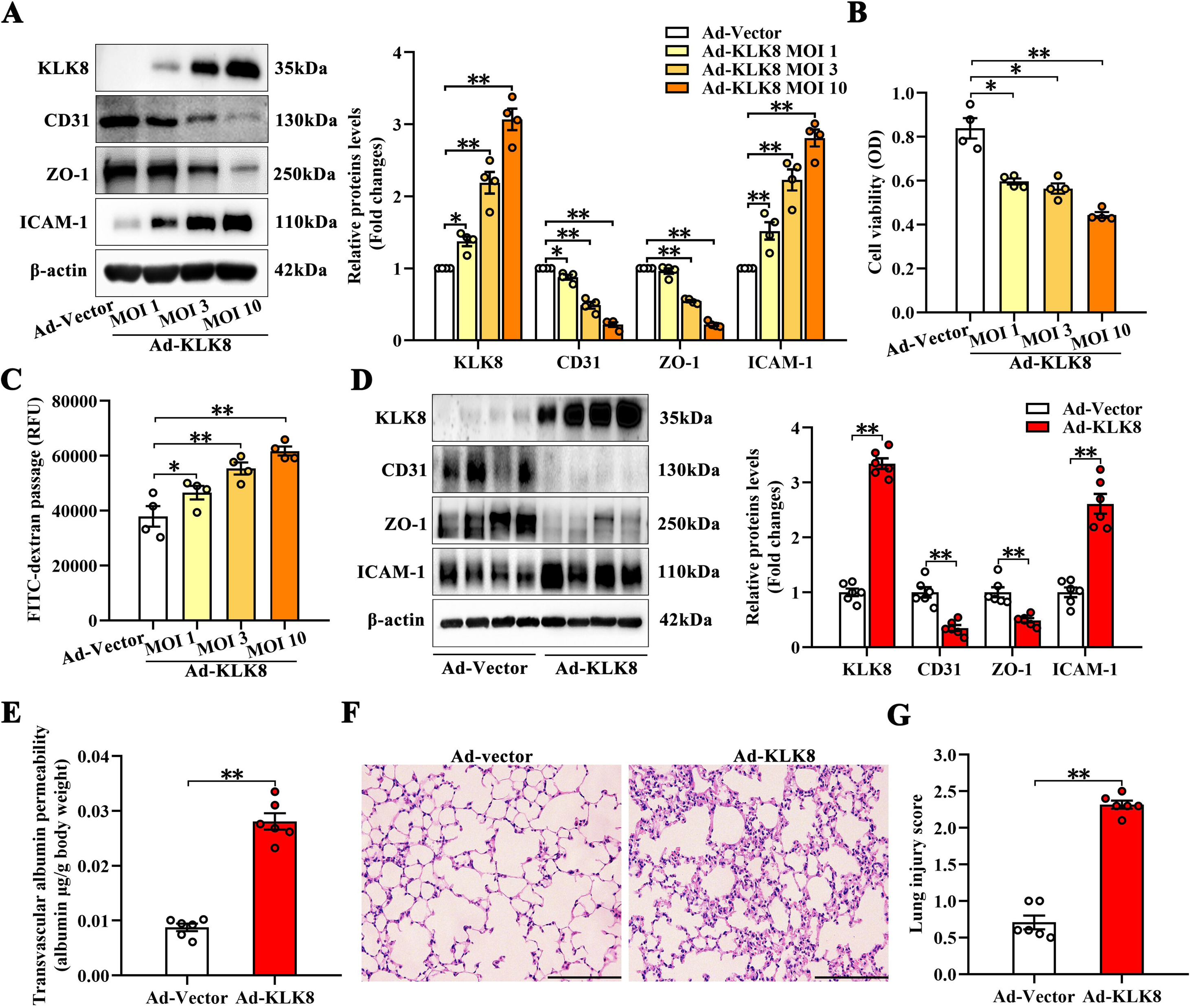
KLK8 overexpression leads to endothelial hyperpermeability and acute lung injury. **A-C**, MLVECs were infected with increasing doses of KLK8 adenovirus (Ad-KLK8) at a multiplicity of infection (MOI) of 1, 3, or 10 for 48 h (n=4). **A**, Protein levels of KLK8, CD31, ZO-1, and ICAM-1 in MLVECs. Corresponding histograms were shown on the right panel of the representative protein bands. **B**, MTT assay demonstrated that Ad-KLK8 dose-dependently decreased cell viability in MLVECs. **C**, Endothelial permeability determined by 10-kDa FITC–dextran flux assay demonstrated that Ad-KLK8 dose-dependently increased endothelial permeability of MLVECs. **D-G**, Mice were exposed to intratracheal instillation of Ad-Vector or Ad-KLK8 for 48 hours (n=6). **D**, Protein levels of KLK8, CD31, ZO-1, and ICAM-1 in lung tissues. Corresponding histograms were shown on the right panel of the representative protein bands. **E**, Pulmonary vascular permeability was quantified by Evans Blue dye extravasation assay into the lung tissue. **F-G**, Lung histopathological evaluation using hematoxylin and eosin staining. **F,** Original magnification, × 200. Scale bars correspond to 100 μm. **G**, The severity of lung injury was scored by two pathologists blinded to group allocation. Statistical analyses were performed using an unpaired, two-tailed Student’s t-test (D, E, G) or one-way ANOVA with Student-Newman-Keuls post hoc test (A-C). Data are presented as the mean ± SEM.. * p<0.05, ** p<0.01.

To assess the effects of KLK8 overexpression *in vivo*, we employed intratracheal transfection of Ad-KLK8. As expected, intra-pulmonary Ad-KLK8 transfection significantly enhanced KLK8 expression in the lung (Figure 2D). Intra-pulmonary KLK8 overexpression resulted in decreased CD31/ZO-1 protein levels and increased ICAM-1 expression (Figure 2D). Measurement of Evans blue dye leakage demonstrated that intra-pulmonary Ad-KLK8 transfection significantly increased pulmonary vascular permeability (Figure 2E). Additionally, histological analysis revealed that mice transfected with intra-pulmonary Ad-KLK8 exhibited significant lung injury, characterized by leukocyte recruitment, diffuse interstitial edema, alveolar thickening, and a profound reduction in alveolar air space, with a lung injury score of 2.32 ± 0.05 (Figure 2F&G).

Taken together, these *in vitro* and *in vivo* studies indicate that overexpression of KLK8 leads to hyperpermeability of the endothelial barrier, ultimately resulting in acute lung injury.

### 2.3 KLK8 upregulation contributes to endothelial hyperpermeability, acute lung injury and mortality induced by systemic LPS administration

As LPS treatment significantly induced KLK8 expression in pulmonary vascular endothelial cells both *in vitro* and *in vivo* (Figure 1), we proceeded to investigate whether the upregulation of KLK8 contributed to LPS-induced endothelial hyperpermeability *in vitro*. Using KLK8 siRNA, we achieved an approximate 80% decrease in KLK8 expression in MLVECs (supplemental Figure S2A). The results presented in Figure 3A-C demonstrated that KLK8 siRNA significantly attenuated LPS-induced endothelial damage, as evidenced by increased CD31/ZO-1 protein levels and cell viability, as well as decreased ICAM-1 expression and FITC-dextran leakage. These findings suggest that LPS may promote endothelial barrier dysfunction through a KLK8-dependent pathway in MLVECs.

**Figure 3.**
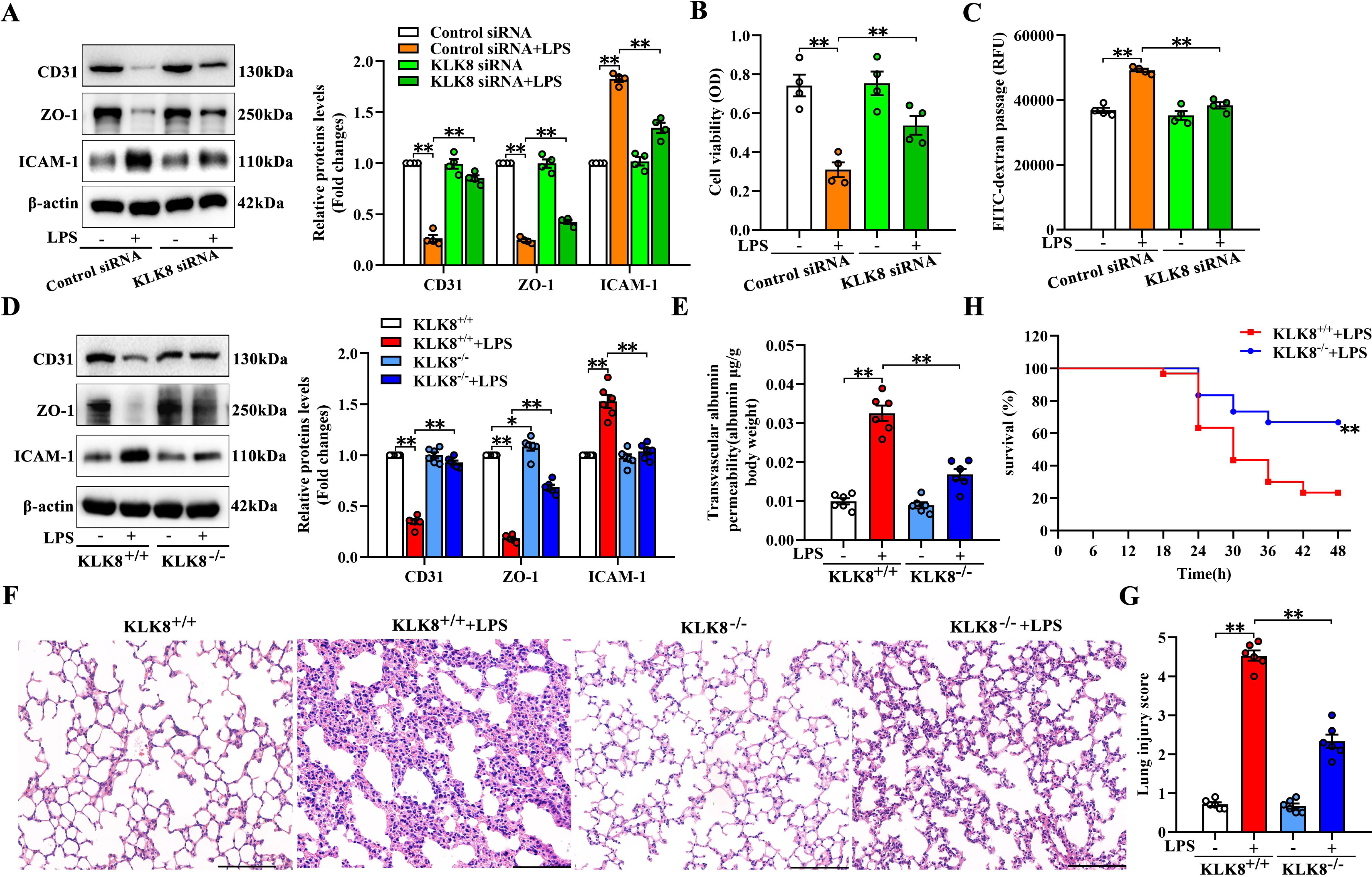
Upregulation of KLK8 contributes to endothelial hyperpermeability, acute lung injury, and mortality induced by systemic LPS administration. **A-C**, MLVECs were transfected with Control siRNA or siRNA targeting KLK8 for 48 hours. Cells were then treated with or without LPS (1,000 ng/ml) for another 24 hours (n=4). **A**, Protein levels of CD31, ZO-1, and ICAM-1 in MLVECs. Corresponding histograms were shown on the right panel of the representative protein bands. **B**, MTT assay showed that KLK8 siRNA attenuated LPS-induced endothelial cell injury. **C**, Endothelial permeability determined by 10-kDa FITC–dextran flux assay showed that KLK8 siRNA attenuated LPS-induced endothelial cell hyperpermeability. **D-G**, KLK8-deficient (KLK8^-/-^) and control (KLK8^+/+^) mice were intraperitoneally injected with 30 mg/kg LPS (n=6). **D**, Protein levels of CD31, ZO-1, and ICAM-1 in lung tissues were examined 24 hours after LPS administration. Corresponding histograms were shown on the right panel of the representative protein bands. **E**, Pulmonary vascular permeability was quantified 24 hours after LPS administration by evans blue dye extravasation assay into the lung tissue. **F-G**, Lung tissue sections were collected 48 hours after LPS administration. Histopathological evaluation were performed using hematoxylin and eosin staining. **F**, Original magnification, × 200. Scale bars correspond to 100 μm. **G**, The severity of lung injury was scored by two pathologists blinded to group allocation. **H**, Kaplan–Meier survival curves for KLK8^+/+^ mice (n =30) and KLK8^-/-^mice (n=30) intraperitoneally injected with 30 mg/kg LPS. Statistical analyses were performed using one-way ANOVA with Student-Newman-Keuls post hoc test (A-E, G) or Kaplan–Meier test (H). Data are presented as the mean ± SEM. * p<0.05, ** p<0.01.

To investigate the impact of KLK8 deficiency on endotoxemia-associated endothelial barrier dysfunction in vivo, we generated global KLK8 deletion by crossing KLK8^f/f^ mice with EIIa cre(+) mice. As expected, pulmonary KLK8 expression was significantly decreased in KLK8-deficient mice compared to wild-type mice (supplemental Figure S2B). KLK8 deficiency reversed the LPS-induced decreases in CD31 and ZO-1 expression (Figure 3D). Additionally, the LPS-induced increases in ICAM-1 expression, Evans blue dye leakage, and lung injury score were profoundly improved in KLK8-deficient mice (Figure 3D-G). Notably, survival studies also revealed that KLK8 deficiency significantly decreased mortality in response to endotoxemia compared to control mice (Figure 3H). These findings indicate that the upregulation of KLK8 contributes to LPS-induced endothelial hyperpermeability, acute lung injury, and mortality.

### 2.4 KLK8 inhibition attenuates endothelial hyperpermeability, acute lung injury and mortality induced by systemic LPS administration or CLP-induced sepsis

To confirm the crucial role of KLK8 in sepsis-associated pulmonary endothelial damage, we further observed the effect of the anti-KLK8 neutralizing antibody on endothelial hyperpermeability, acute lung injury and mortality induced by systemic LPS administration or CLP-induced sepsis. As depicted in Figure 4A-D, the anti-KLK8 blocking antibody effectively reduced LPS-induced endothelial hyperpermeability and acute lung injury, as evidenced by increased CD31 and ZO-1 expression, as well as decreased ICAM-1 expression, Evans blue dye leakage, and lung injury score. The protective effects of the anti-KLK8 blocking antibody in CLP-induced septic mice (Figure 5A-D) also mirrored the results observed in LPS-treated mice. Furthermore, mice treated with the anti-KLK8 neutralizing antibody exhibited a higher survival rate compared to control IgG-treated mice exposed to LPS (Figure 4E) or CLP (Figure 5E).

**Figure 4.**
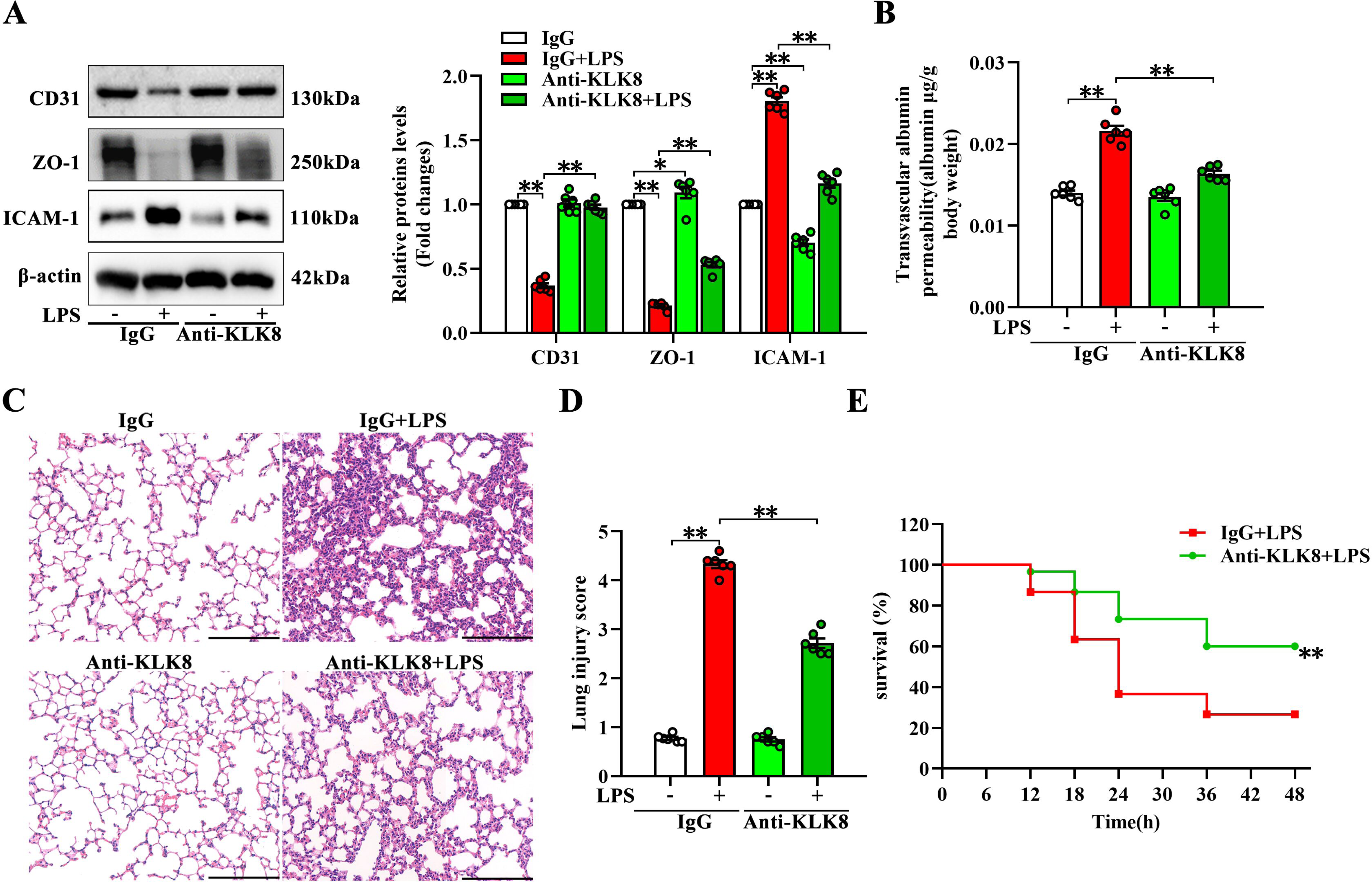
KLK8 inhibition attenuates endothelial hyperpermeability, acute lung injury, and mortality induced by systemic LPS administration. **A-D,** Mice were treated with anti-KLK8 neutralizing antibodies or IgG isotype via the tail vein. Thirty minutes later, mice were injected intraperitoneally with LPS at 30 mg/kg or saline control (n=6). **A**, Protein levels of CD31, ZO-1, and ICAM-1 in lung tissues were examined 24 hours after LPS administration. Corresponding histograms were shown on the right panel of the representative protein bands. **B**, Pulmonary vascular permeability was quantified 24 hours after LPS administration by evans blue dye extravasation assay into the lung tissue. **C-D,** Lung tissue sections were collected 48 hours after LPS administration. Histopathological evaluation were performed using hematoxylin and eosin staining. **C**, Original magnification, × 200. Scale bars correspond to 100 μm. **D**, The severity of lung injury was scored by two pathologists blinded to group allocation. **E**, Kaplan–Meier survival curves for anti-KLK8 neutralizing antibodies-treated (n=30) or IgG isotype-treated (n =30) mice intraperitoneally injected with 30 mg/kg LPS. Statistical analyses were performed using one-way ANOVA with Student-Newman-Keuls post hoc test (A, B, D) or Kaplan–Meier test (E). Data are presented as the mean ± SEM. * p<0.05, ** p<0.01.

**Figure 5.**
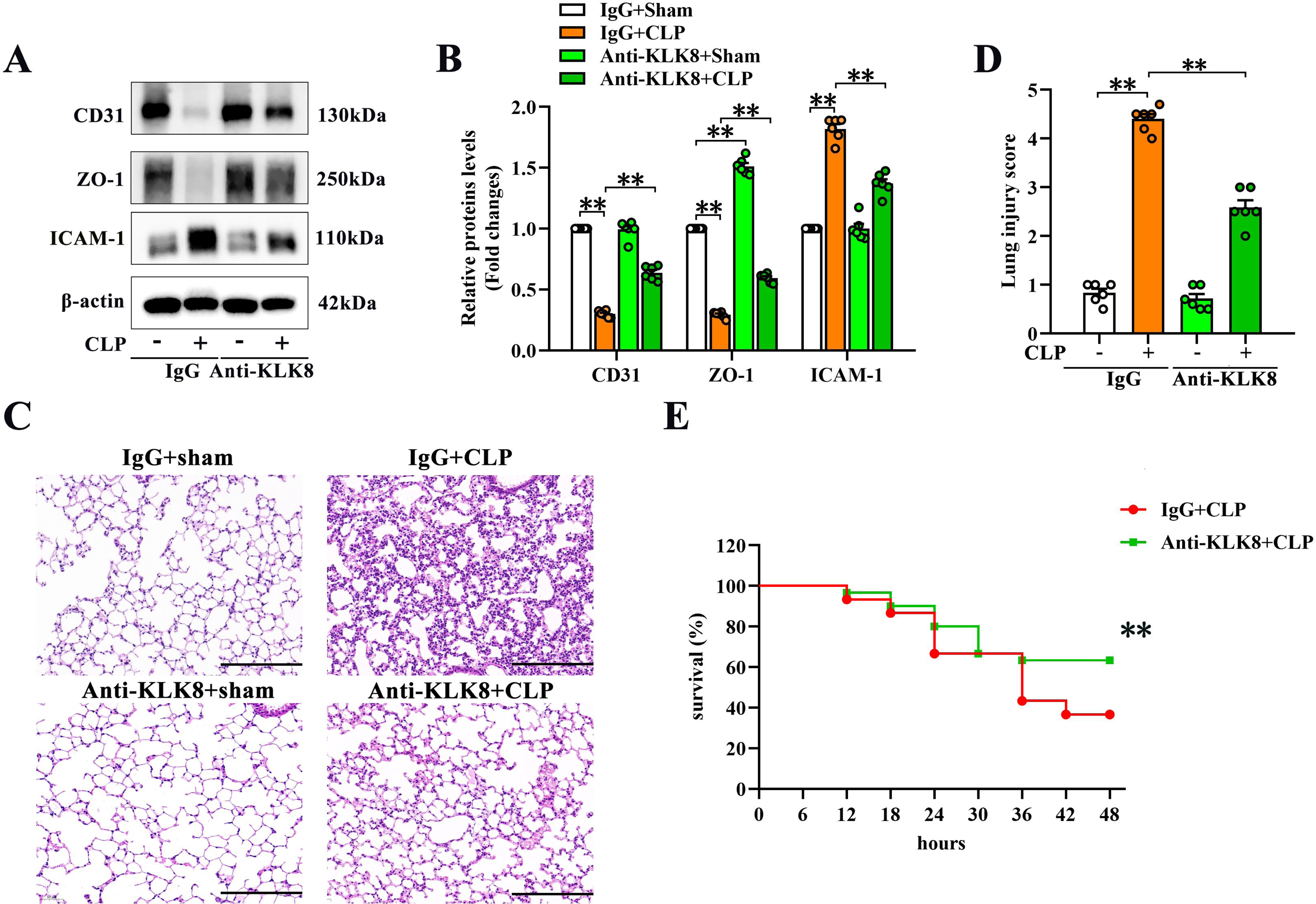
KLK8 inhibition attenuates endothelial injury, acute lung injury, and mortality induced by CLP-induced sepsis. **A-D,** Mice were treated with anti-KLK8 neutralizing antibodies or IgG isotype via the tail vein. Thirty minutes later, mice were subjected to CLP or Sham surgery (n=6). **A-B**, Protein levels of CD31, ZO-1, and ICAM-1 in lung tissues were examined 24 hours after CLP or Sham surgery. **B** showed corresponding histograms of the protein bands. **C-D,** Lung tissue sections were collected 48 hours after CLP or Sham surgery. Histopathological evaluation were performed using hematoxylin and eosin staining. **C**, Original magnification, × 200. Scale bars correspond to 100 μm. **D**, The severity of lung injury was scored by two pathologists blinded to group allocation. **E**, Kaplan–Meier survival curves for anti-KLK8 neutralizing antibodies-treated (n=30) or IgG isotype-treated (n =30) mice subjected to CLP or Sham surgery. Statistical analyses were performed using one-way ANOVA with Student-Newman-Keuls post hoc test (B, D) or Kaplan–Meier test (E). Data are presented as the mean ± SEM. * p<0.05, ** p<0.01.

A previous study has demonstrated that continuous intraventricular administration of the anti-KLK8 blocking antibody for 4 weeks exhibited anxiolytic effects in wild-type mice, as evidenced by increased exploratory behavior in the open field tests (Herring et al., 2016). We then observed the effects of the anti-KLK8 antibody on behavioral performance in mice. As shown in supplemental Figure S3, systemic LPS administration resulted in significant decreases in the distance or time spent in the central area, the ratio of the distance or time in the central area compared to that in the periphery, and the number of crossing squares. These effects were modestly attenuated by the injection of the anti-KLK8 neutralizing antibody. Additionally, it was found that the anti-KLK8 antibody itself had no significant effects on behavioral parameters in the open field tests (supplemental Figure S3).

### 2.5 KLK8 induces endothelial barrier dysfunction by suppressing the VE-cadherin/Akt/FOXM1 signaling pathway-mediated endothelial cell proliferation

To investigate the mechanisms underlying KLK8-induced endothelial injury, we performed RNA-Seq analysis on MLVECs treated with Ad-KLK8 compared to those treated with Ad-Vector (n=4 per group). We identified 996 up-regulated and 2368 down-regulated DEGs in Ad-KLK8-treated MLVECs compared to Ad-Vector-treated MLVECs (|LogFC| ≥ 1, false discovery rate < 0.05, Figure 6A-B). GSEA was then performed to determine the enriched biological pathways in KLK8-overexpressed endothelial cells. Following Hallmark, Kyoto Encyclopedia of Genes and Genomes (KEGG), Gene Ontology (GO), and Reactome pathway analysis of GSEA, gene sets associated with lactate metabolic process, intrinsic apoptotic signaling pathway, and FGFR1 ligand binding and activation were significantly enriched in the Ad-KLK8-treated MLVECs (Supplemental Figure S4). Conversely, gene sets associated with cell cycle, DNA repair, vascular wound healing, establishment of endothelial barrier and endothelial cell proliferation were significantly enriched in the Ad-Vector-treated cells (Figure 6C and Supplemental Figure S5-6).

**Figure 6.**
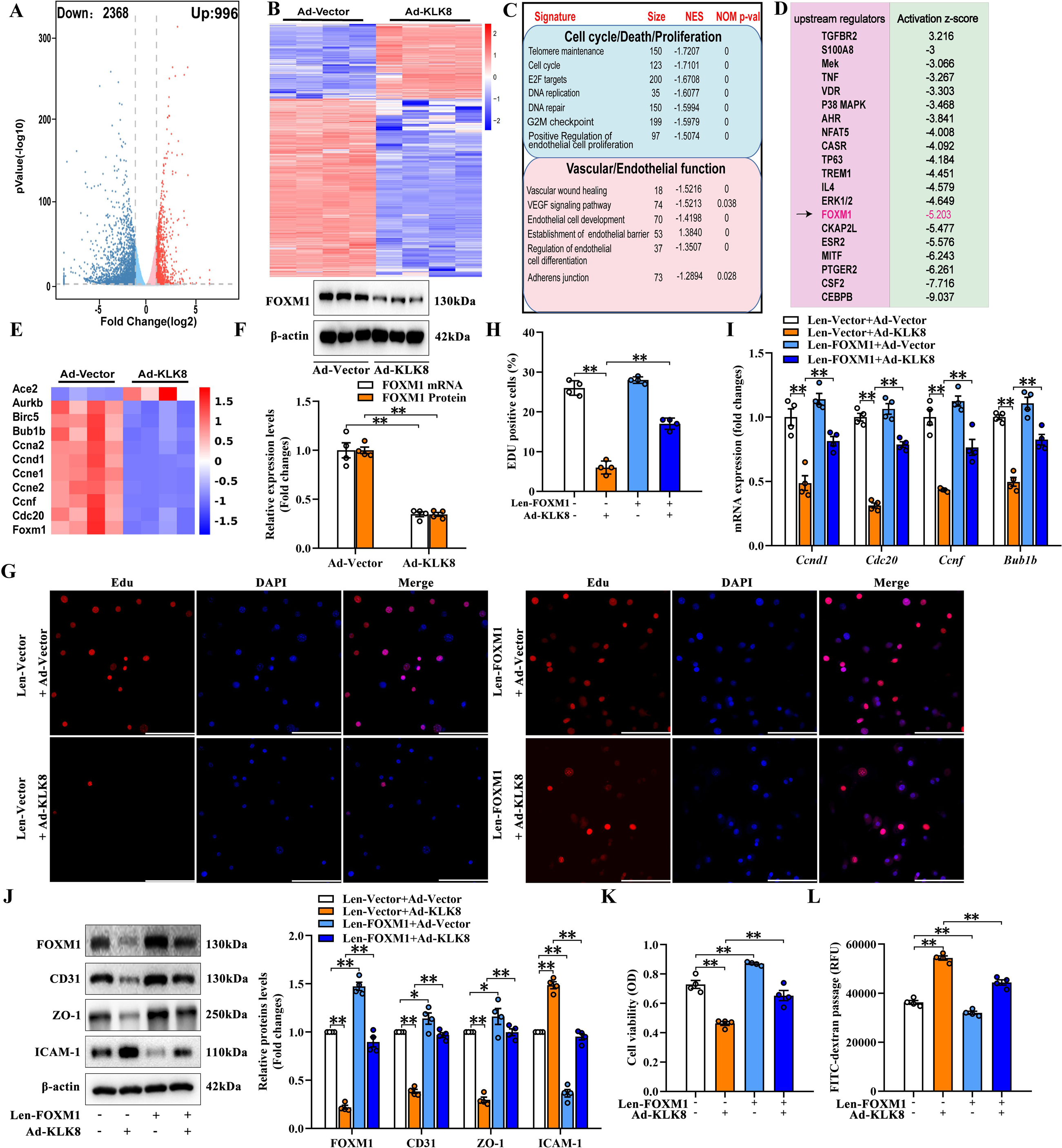
KLK8 induces endothelial barrier dysfunction by suppressing FOXM1-mediated endothelial cell proliferation. **A-E**, MLVECs were infected with Ad-Vector or Ad-KLK8 at a multiplicity of infection (MOI) of 3 for 48 hours. Dysregulated genes were analyzed by RNA-Seq. **A**, Volcano plots showing differentially expressed genes (DEGs) in KLK8-overexpressed MLVECs. Red reflects upregulated and blue downregulated genes. **B**, Heat map of DEGs, red indicating upregulated and blue downregulated genes. **C**, Molecular signatures depicting the dysregulated pathways, along with the number of genes in each pathway, normalized enrichment score (NES), and normalized (NOM) p value. **D**, Ingenuity Pathway Analysis (IPA) of the DEGs predicts the upstream regulators responsible for alterations of the gene expression profile in KLK8-overexpressed MLVECs. **E**, Heat map of FOXM1 and its target molecules in dataset of IPA, red indicating upregulated and blue downregulated genes. **F**, The mRNA and protein levels of FOXM1 in Ad-KLK8 treated MLVECs, corresponding histograms were presented below the protein bands. **G-L**, MLVECs were infected with Lentivirus vector (Len-Vector) or FOXM1 lentivirus (Len-FOXM1) at a MOI of 10 for 30 min, and then treated with Ad-Vector or Ad-KLK8 at a MOI of 3 for 48 hours. **G-H**, After treatment with Len-FOXM1 with or without Ad-KLK8, the cells were incubated in medium containing 10 μM EdU for 2 h. Cell proliferation was determined using Click™ EdU-594 (red) Kit. The nucleus was stained with DAPI (blue). **H**, Cell proliferation was quantified by showing the percentage of EdU-positive nuclei in DAPI-positive nuclei. **I**, The mRNA levels of FOXM1 target molecules *Ccnd1*, *Cdc20*, *Ccnf*, and *Bub1b*. **J**, Protein levels of FOXM1, CD31, ZO-1, and ICAM-1 in MLVECs. Corresponding histograms were shown on the right panel of the representative protein bands. **K**, MTT assay showed that FOXM1 overexpression attenuated KLK8 overexpression-induced endothelial cell injury. **L**, Endothelial permeability determined by 10-kDa FITC–dextran flux assay showed that FOXM1 overexpression markedly decreased KLK8 overexpression-induced endothelial cell hyperpermeability. Statistical analyses were performed using an unpaired, two-tailed Student’s t-test (F) or one-way ANOVA with Student-Newman-Keuls post hoc test (I, J, K, L). Data are presented as the mean ± SEM (n=4). * p<0.05; ** p<0.01.

Furthermore, Ingenuity Pathway Analysis was performed to predict the upstream regulators responsible for the altered gene expression profile in KLK8-overexpressed endothelial cells. As shown in Figure 6D and supplemental Table 2, a total of 26 upstream regulators were identified to be either activated (z-score > 2, p value of overlap < 0.01) or inhibited (z-score < -2, p value of overlap < 0.01). These regulators are involved in various biological processes, including cell cycle regulation, kinase signaling, transcriptional regulation, and immune system responses. Notably, FOXM1, a transcription factor essential for endothelial development and regeneration (Zhao et al., 2006; Huang et al., 2016), was predicted to be highly inhibited (z-score -5.203) and was profoundly downregulated (LogFC -2.38, p < 0.001) in Ad-KLK8-treated MLVECs (Figure 6E). Analysis of the RNA-seq data revealed that FOXM1 targets involved in cell proliferation and cell cycle progression were significantly downregulated in KLK8-overexpressed endothelial cells (e.g., *Aurkb, Birc5, Bub1b, Ccna2, Ccnd1, Ccne1, Ccne2, Ccnf,* and *Cdc20*) (Figure 6E).

Validation through quantitative RT-PCR and western blotting confirmed the downregulation of FOXM1 in MLVECs treated with Ad-KLK8 (Figure 6F). We then assessed the effect of Ad-KLK8-induced FOXM1 suppression on endothelial cell proliferation using EdU incorporation. Ad-KLK8-treated MLVECs exhibited a significant decrease in cell proliferation compared to Ad-Vector-treated MLVECs, which was restored by lentivirus-mediated FOXM1 overexpression (Figure 6G&H). To further understand the basis of inhibited cell proliferation in KLK8-overexpressed endothelial cells, we examined the mRNA levels of FOXM1 target genes responsible for cell proliferation using QRT-PCR analysis. The mRNA levels of *Ccnd1, Cdc20, Ccnf,* and *Bub1b* were found to be downregulated in Ad-KLK8-treated MLVECs compared to Ad-Vector-treated MLVECs (Figure 6I). However, FOXM1 overexpression significantly increased the mRNA levels of these genes in MLVECs. Additionally, FOXM1 overexpression markedly attenuated KLK8 overexpression-induced endothelial damage, as evidenced by increased CD31/ZO-1 protein levels and cell viability, as well as decreased ICAM-1 expression and FITC-dextran leakage (Figure 6J-L). Collectively, our results indicate that KLK8 may induce endothelial barrier dysfunction by suppressing FOXM1-mediated endothelial cell proliferation.

To further understand the mechanisms underlying KLK8-induced downregulation of FOXM1, we conducted single-gene GSEA using Hallmark gene sets based on RNA-seq data from KLK8-overexpressed MLVECs. Interestingly, we observed a positive correlation between FOXM1 expression and gene sets associated with the PI3K-AKT-MTOR and MTORC1 signaling pathways (p < 0.01, Figure 7A and supplemental Figure S7A), both of which have been previously reported to modulate FOXM1 expression (Zhang et al., 2017; Zarrouki et al., 2014). We then investigated whether the PI3K-Akt or MTORC1 pathways were involved in the KLK8-induced downregulation of FOXM1. As shown in Figure 7B, treatment with Ad-KLK8 significantly decreased Akt phosphorylation, while having no significant effects on the phosphorylation of mTOR and its downstream signaling proteins p70 S6K (supplemental Figure S7B). Additionally, we found that SC79, a small-molecule Akt activator, not only stimulated basal Akt phosphorylation and FOXM1 expression but also completely reversed the KLK8-induced downregulation of FOXM1 in endothelial cells (Figure 7C). On the other hand, the mTOR agonist MHY1485 did not affect FOXM1 expression, although it increased the phosphorylation of mTOR and p70 S6K (supplemental Figure S7C). These results suggest that KLK8 may suppress FOXM1 expression through the inactivation of the Akt signaling pathway.

**Figure 7.**
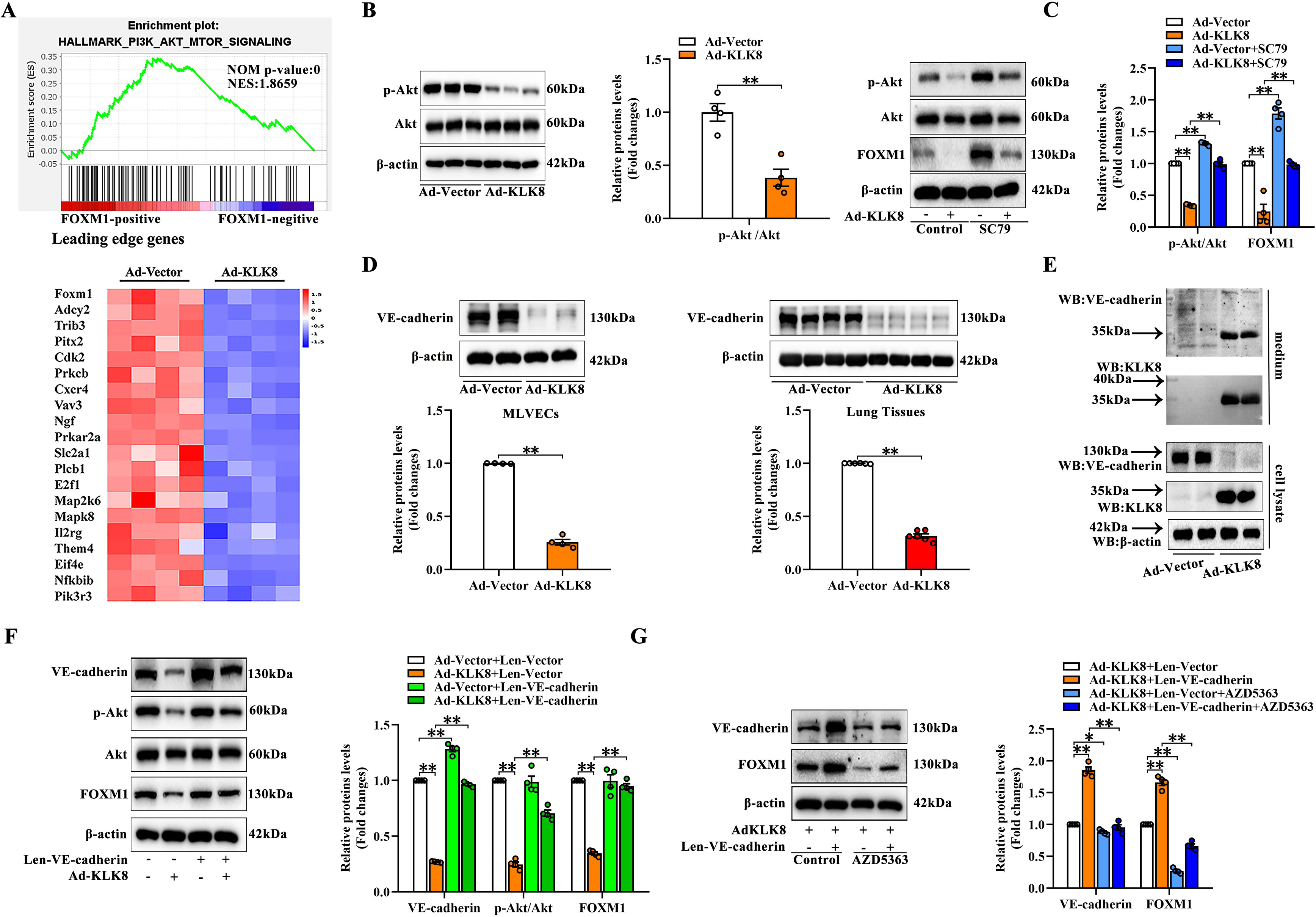
KLK8 suppresses FOXM1 expression by inactivating the VE-cadherin/Akt signaling pathway. **A**, Gene Set Enrichment Analysis (GSEA) plot of genes regulated by PI3K-AKT-MTOR signaling pathway, showing that activation of the pathway was positively related with FOXM1 expression in RNA-seq data of Ad-KLK8 treated MLVECs. Heat map of FOXM1 and the dysregulated target genes of PI3K-AKT-MTOR signaling pathway in KLK8-overexpressed MLVECs were presented below the GSEA plot. Red reflects upregulated and blue downregulated genes. **B**, Protein levels of phosphorylated Akt and Akt in MLVECs were examined 48 hours after infection of Ad-Vector or Ad-KLK8 at MOI of 3. Corresponding histograms were shown on the right panel of the representative protein bands (n=4). **C**, Protein levels of phosphorylated Akt, Akt, and FOXM1 in MLVECs were examined 48 hours after infection of Ad-Vector or Ad-KLK8 at MOI of 3 in the presence or absence of Akt activator SC79 (10 μM). Corresponding histograms were shown on the right panel of the representative protein bands (n=4). **D**, Protein levels of VE-cadherin in MLVECs (n=4, left panel) or lung tissues (n=6, right panel) exposed to Ad-KLK8 for 48 hours. Corresponding histograms were shown below the protein bands. **E**, MLVECs were treated with Ad-vector or Ad-KLK8 (MOI of 3) in serum-free medium for 48 h. Westen blot analysis exhibited the appearance of a ∼30 kDa N-terminal VE-cadherin fragment in the culture medium. **F**, MLVECs were infected with Lentivirus vector (Len-Vector) or VE-cadherin lentivirus (Len-VE-cadherin) at a MOI of 10 for 30 min, and then treated with Ad-Vector or Ad-KLK8 at a MOI of 3. Protein levels of VE-cadherin, FOXM1, phosphorylated Akt and Akt in MLVECs were examined 48 hours after infection of Ad-Vector or Ad-KLK8. Corresponding histograms were shown on the right panel of the representative protein bands (n=4). **G**, Protein levels of VE-cadherin and FOXM1 in MLVECs were examined 48 hours after infection of Ad-KLK8 at MOI of 3 in the presence or absence of Len-VE-cadherin or Akt inhibitor AZD5363 (10 μM). Corresponding histograms were shown on the right panel of the representative protein bands (n=4). Statistical analyses were performed using an unpaired, two-tailed Student’s t-test (B, D) or one-way ANOVA with Student-Newman-Keuls post hoc test (C, F, G). Data are presented as the mean ± SEM. * p<0.05; ** p<0.01.

Recently, Du et al. reported that KLK8 cleaves VE-cadherin, thereby promoting EndMT in human coronary artery endothelial cells (Du et al., 2021). Cleavage of VE-cadherin is known to inactivate the Akt signaling pathway in endothelial cells (Carmeliet et al., 1999). Therefore, we investigated whether the cleavage of VE-cadherin was involved in the KLK8-induced inactivation of Akt. As shown in Figure 7D, overexpression of KLK8 downregulated VE-cadherin in both MLVECs and lung tissues. We then examined the cell culture medium of Ad-KLK8-treated MLVECs using western blot analysis with an antibody recognizing the N-terminus of VE-cadherin. We observed a new ∼30 kDa fragment of VE-cadherin in the culture medium of Ad-KLK8-treated endothelial cells (Figure 7E), which is consistent with KLK8 cleaving VE-cadherin in the extracellular cadherin domain, as reported in coronary artery endothelial cells (Du et al., 2021). Furthermore, lentivirus-mediated overexpression of VE-cadherin significantly increased Akt phosphorylation and FOXM1 expression in KLK8-overexpressed MLVECs (Figure 7F). Moreover, the stimulatory effect of VE-cadherin overexpression on FOXM1 expression was largely blocked by the Akt inhibitor AZD5363 (Figure 7G). These findings indicate that KLK8 suppresses FOXM1 expression by inactivating the VE-cadherin/Akt signaling pathway.

### 2.6 KLK8 deficiency or blockade rescued VE-cadherin/Akt/FOXM1 signaling pathway, thus promoting endothelium regeneration

As described above, KLK8 deficiency or blockade significantly attenuated endothelial barrier dysfunction induced by systemic LPS administration or CLP-induced sepsis (Figure 3-5). We then investigated whether KLK8 deficiency or blockade could restore the VE-cadherin/Akt/FOXM1 signaling pathway and promote endothelial regeneration after injury. We observed that both systemic LPS administration and CLP-induced sepsis led to decreased levels of VE-cadherin, Akt phosphorylation, and FOXM1, which were restored in mice treated with anti-KLK8 antibody (Figure 8A-B and supplemental Figure S8). Furthermore, the LPS-induced decreases in VE-cadherin, Akt phosphorylation, and FOXM1 were also reversed in KLK8-deficient mice (supplemental Figure S9A).

**Figure 8.**
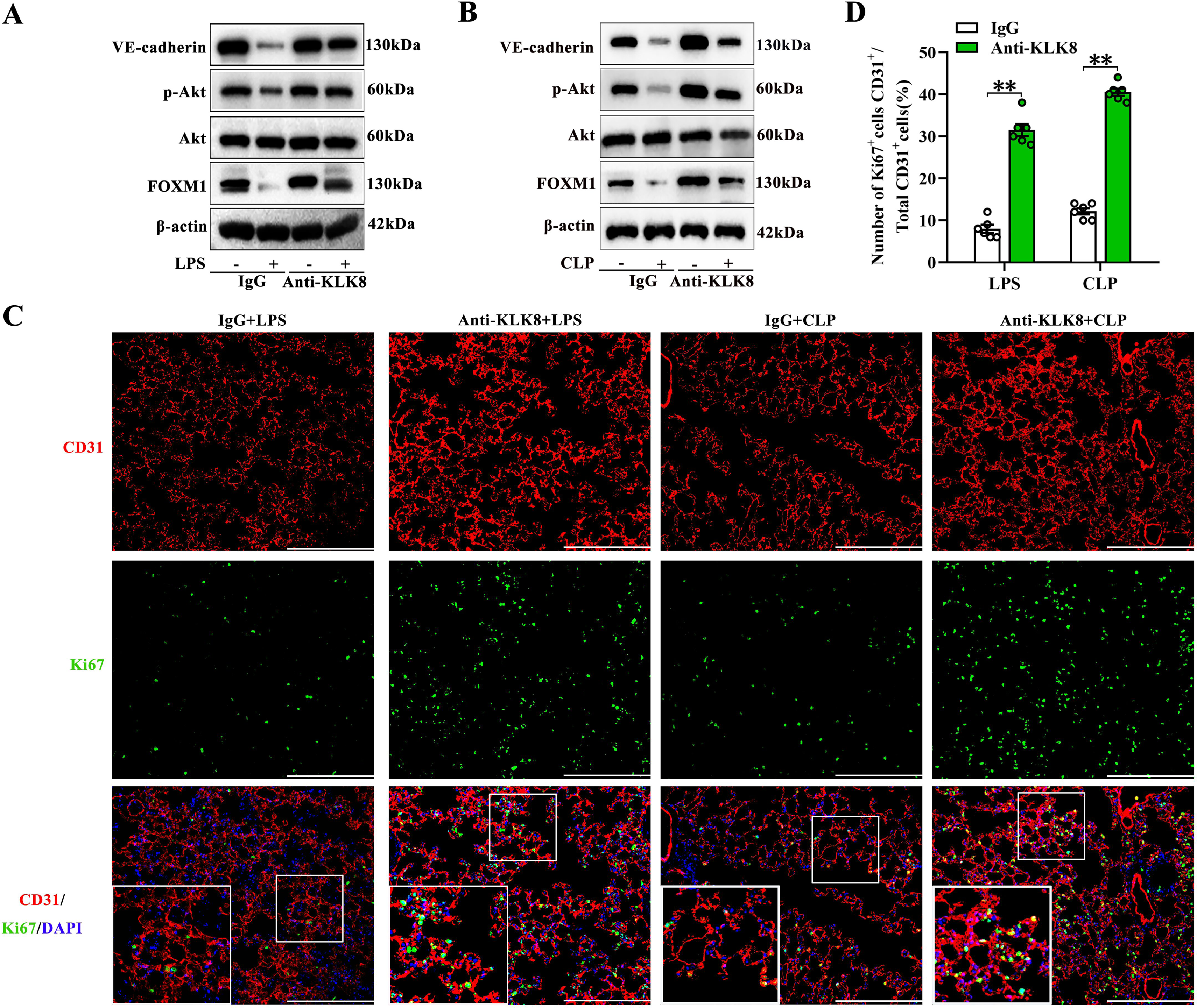
KLK8 blockade rescues VE-cadherin/Akt/FOXM1 signaling pathway, thus promoting endothelium regeneration. Mice were treated with anti-KLK8 neutralizing antibodies or IgG isotype via the tail vein. Thirty minutes later, mice were injected intraperitoneally with LPS at 30 mg/kg or saline control, or subjected to CLP or sham surgery. **A-B**, Protein levels of VE-cadherin, phosphorylated Akt, Akt, and FOXM1 in lung tissues were examined 24 hours after LPS administration (A) or 24 hours after CLP/sham surgery (B). **C-D**, Lung tissue sections were collected 48 hours after LPS administration or CLP/sham surgery. Lung sections were stained with fluorophore-labeled antibodies against endothelial cell marker CD31 (red) and cell proliferation marker Ki-67 (green). Nuclei were counterstained with DAPI (blue). Scale bars correspond to 100 μm. **D**, Quantification of the percentage of Ki67^+^/CD31^+^ cells in total CD31^+^ cells in lung tissue sections obtained from anti-KLK8/IgG-treated mice subjected to LPS or CLP-induced sepsis. Statistical analyses were performed using an unpaired, two-tailed Student’s t-test. Data are presented as the mean ± SEM (n=6). ** p<0.01.

To assess the effect of KLK8 deficiency or blockade on endothelial cell proliferation, we performed double immunofluorescence staining against CD31 and the cell proliferation marker Ki67 in lung sections. Control mouse lungs showed endothelial cell proliferation 48 hours after LPS administration or CLP (Figure 8C and D). Interestingly, we observed a significant increase in the colocalization of CD31/Ki67 in lung tissues of mice treated with anti-KLK8 antibody compared to IgG-treated mice (Figure 8C and D). Similarly, LPS-treated KLK8-deficient mice exhibited an increased number of CD31^+^/Ki67^+^ cells in lung tissue sections compared to LPS-treated KLK^+/+^ mice (supplemental Figure S9B and C). These findings collectively demonstrate that KLK8 deficiency or blockade restores the VE-cadherin/Akt/FOXM1 signaling pathway, thereby promoting endothelial regeneration.

## 3. Discussion

Endothelial dysfunction and capillary hyperpermeability play crucial roles in sepsis-associated acute respiratory distress syndrome (Joffre et al., 2020; Dolmatova et al., 2021; De Backer et al., 2021). Pulmonary endothelial cells are particularly susceptible to injury from inflammatory stimuli such as LPS and bacterial infections (Vassiliou et al., 2020; Ince et al., 2016). This study has uncovered a novel function of KLK8 in mediating lung endothelial barrier dysfunction during endotoxemia and CLP-induced sepsis. The degradation of VE-cadherin induced by KLK8 and the subsequent inactivation of the Akt/FOXM1 pathway impair endothelial regeneration, contributing to pulmonary vascular endothelial barrier dysfunction during sepsis (Figure 9).

**Figure 9.**
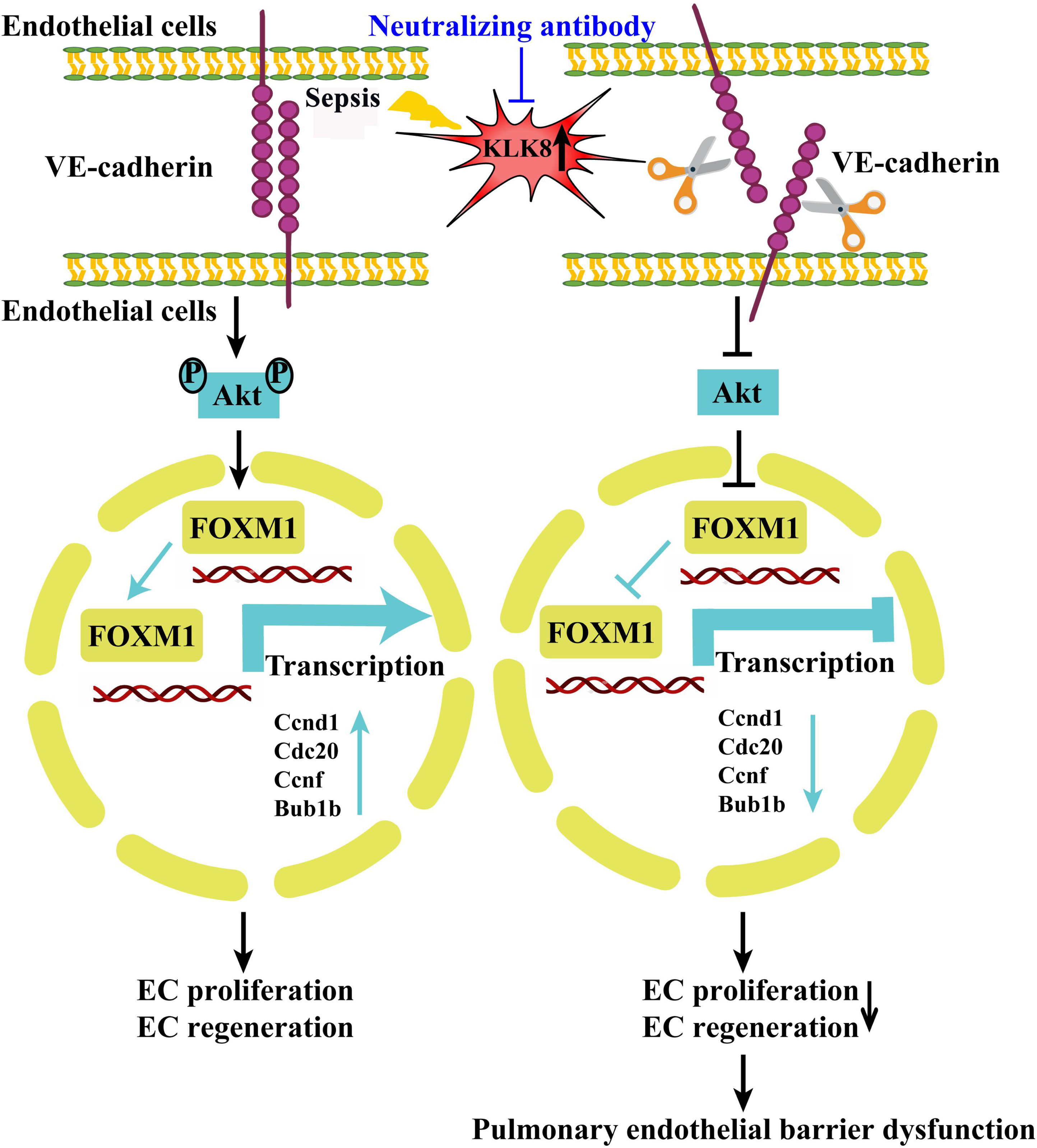
Schematic diagram of the mechanism by which upregulated KLK8 contributes to the development of sepsis-induced pulmonary endothelial barrier dysfunction. KLK8 is upregulated in pulmonary vascular endothelial cells during sepsis. As a secreted serine protease, KLK8 cleavages the extracellular domain of VE-cadherin and inactivates Akt/FOXM1 signaling pathway. KLK8-induced inactivation of VE-cadherin/Akt/FOXM1 pathway inhibits the expression of FOXM1 target genes involved in cell proliferation and cell cycle progression, which finally suppresses FOXM1-mediated endothelial cell (EC) proliferation and regeneration and results in pulmonary endothelial barrier dysfunction. KLK8 inhibition by anti-KLK8 neutralizing antibodies may represent a novel therapeutic strategy for restoring the microvascular barrier integrity during sepsis.

KLK8 is normally expressed at low levels in lung tissues but becomes upregulated in pulmonary vascular endothelial cells in endotoxemic and septic mice, suggesting it as a potential therapeutic target for inflammation-associated pulmonary endothelial dysfunction. According to previous literature (Lenga Ma Bonda et al., 2018), KLK8 expression in normal tissues is limited to basal cells of bronchi, goblet cells, and submucosal glands. Consistently, we observed several KLK8+ cells in the perivascular area of lung tissues from control, endotoxemic, and septic mice. However, our results also revealed strong KLK8 staining within the microvascular endothelial cells in endotoxemic and septic lungs, in contrast to low staining for KLK8 in control lung tissue sections.

Another member of the KLK family, KLK1 (also known as tissue kallikrein), is implicated in the pathogenesis of sepsis. Ran et al. recently reported a significant increase in plasma KLK1 levels in septic patients compared to non-sepsis controls, with levels positively associated with the severity and mortality of sepsis (Ran et al., 2021). In in vitro studies using human umbilical vein endothelial cells, they demonstrated that KLK1 treatment exacerbates, while the KLK1 inhibitor kallistatin significantly mitigates LPS-induced endothelial hyperpermeability (Ran et al., 2021). However, the involvement of other KLKs in sepsis-associated endothelial dysfunction remains largely unknown. In our study, we found that KLK8, but not KLK1, was the most highly upregulated KLK in endotoxemic lungs and LPS-treated MLVECs, suggesting its primary role in mediating inflammation-induced pulmonary endothelial barrier dysfunction. KLK8 overexpression induced endothelial hyperpermeability both in vitro and in vivo. Conversely, inhibition of KLK8 significantly alleviated endothelial hyperpermeability, acute lung injury, and mortality induced by LPS or CLP-induced sepsis. These findings suggest that KLK8 upregulation contributes to sepsis-induced pulmonary endothelial barrier dysfunction. It is worth noting that KLKs are secreted as proenzymes, which are further processed by other KLKs or proteases to become active KLKs (Prassas et al., 2015). A previous study has identified the ability of KLK8 to activate pro-KLK1 in vitro, but not vice versa (Eissa et al., 2011). The fact that KLK8 acts upstream of KLK1 in the KLK proteolytic cascade suggests that KLK8 may represent a better target for inhibition than KLK1.

Transcription profiling of endothelial cells overexpressing KLK8 revealed that downregulation of FOXM1 plays a central role in mediating the detrimental effects of KLK8 on pulmonary endothelial cells, inhibiting their proliferation. This finding aligns with previous reports on the role of FOXM1 in pulmonary vascular endothelial cell proliferation and repair in response to endotoxemia (Zhao et al., 2006, Zhao et al., 2014), sepsis (Huang et al., 2019, Huang and Zhao, 2012), and hyperoxia (Bolte et al., 2020). However, the mechanisms regulating FOXM1 expression in endothelial cells are not well understood. In this study, we identified Akt as a potential upstream regulator of FOXM1 expression in pulmonary endothelial cells, while mTOR may not be involved. Interestingly, FOXM1 is known to be repressed by p53 in cancer cells (Barsotti and Prives, 2009, Bollu et al., 2020), and there is reciprocal inhibition between Akt and p53 in endothelial cells (Wang et al., 2007, Yokoyama et al., 2014). Previous research has shown that KLK8 induces p53 expression in human coronary artery endothelial cells (Du et al., 2021). We observed that KLK8 inactivated Akt in MLVECs, suggesting a possible feedback regulatory loop between Akt and p53 that contributes to the suppression of FOXM1 expression by KLK8 in endothelial cells. Further investigation is needed to explore this potential regulatory loop.

Endothelial regeneration and the re-establishment of endothelial adherens junctions are crucial for repairing the endothelial barrier and restoring vascular integrity (Lampugnani et al., 2018, Komarova et al., 2017). VE-cadherin, a major component of endothelial adherens junctions, also plays a role in vascular homeostasis by transmitting intracellular signals (Yang and Fan, 2022, Giannotta et al., 2013). The discovery that KLK8 degrades VE-cadherin prompted us to investigate VE-cadherin signaling pathways as mediators of KLK8-induced inhibition of FOXM1 expression. Our study provided clear evidence that lentivirus-mediated overexpression of VE-cadherin significantly increased Akt phosphorylation and FOXM1 expression in KLK8-overexpressed MLVECs. This effect was blocked by an Akt inhibitor. Moreover, inhibition of KLK8 restored the VE-cadherin/Akt/FOXM1 signaling pathway and promoted endothelial regeneration in lung tissues of endotoxemic or CLP-induced septic mice. Considering the pro-proliferative and pro-survival effects of FOXM1 on endothelial cells, our findings suggest that restoration of VE-cadherin by inhibiting KLK8 not only strengthens endothelial adherens junctions but also contributes to endothelial regeneration and repair after vascular injury by stimulating Akt-dependent FOXM1 expression.

It is worth noting that FOXM1 has been extensively studied for its protective properties in inflammation-associated acute lung injury (Zhao et al., 2006, Zhao et al., 2014, Huang et al., 2019, Huang and Zhao, 2012). However, the concept of upregulating FOXM1 expression or activity for therapeutic endothelial regeneration has limitations. FOXM1 activation may have potential side effects, as it promotes tumor growth and angiogenesis in various malignancies (Yao et al., 2018, Castaneda et al., 2022, Wang et al., 2018). In contrast, KLK8 is known to promote the proliferation and invasiveness of cancer cells in the pancreas, stomach, and colon (Hua et al., 2021a, Lim et al., 2020, Hua et al., 2021b). Based on our findings, inhibiting KLK8 may be a more effective therapeutic strategy for enhancing endothelial regeneration. We demonstrated that systemic delivery of anti-KLK8 neutralizing antibodies enhanced endothelial regeneration, reduced endothelial barrier hyperpermeability, and decreased mortality. Therefore, KLK8 inhibition could be a useful approach for restoring endothelial integrity and mitigating sepsis-associated lung injury. It is important to acknowledge the main limitation of this study. Since we only assessed the prophylactic effects of anti-KLK8 antibody treatment in murine models of endotoxemia and CLP-induced sepsis, further studies are needed to determine whether administering anti-KLK8 antibodies after the onset of sepsis is therapeutically beneficial for sepsis-associated endothelial dysfunction and lung injury.

In summary, this study identified a novel role for upregulated KLK8 in the development of sepsis-induced pulmonary endothelial barrier dysfunction. Inhibiting KLK8 significantly alleviated endothelial hyperpermeability, acute lung injury, and mortality induced by LPS or CLP-induced sepsis. Mechanistically, we provided evidence supporting the critical role of KLK8-induced inactivation of the VE-cadherin/Akt/FOXM1 pathway in impairing endothelial regeneration and causing lung vascular leakage in response to LPS or CLP. Therefore, KLK8 inhibition using neutralizing antibodies may represent a promising therapeutic strategy for restoring microvascular barrier integrity during sepsis.

## 4. Material and methods

### 4.1 Murine models of endotoxemia, CLP-induced sepsis, and anti-KLK8 neutralizing antibody treatment

All laboratory mice in this study were housed in a pathogen-free facility at the Animal Research Center of Navy Medical University. The animal studies were conducted in accordance with the Guidelines for the Care and Use of Laboratory Animals published by the NIH (NIH publication No. 85-23, revised 1996), and were approved by the Ethics Committee of Navy Medical University. It is important to note that only male mice were included in all *in vivo* experiments, and both genotype and treatment were blinded during the measurement and analysis stages. The KLK8-flox mouse line was generated at the Shanghai Biomodel Organism Science & Technology Development Co., Ltd (Shanghai, China) using a LoxP targeting system with two LoxP elements flanking exon 1-3 of KLK8, as previously described (Du et al., 2021). To generate global KLK8 knockout mice, KLK8-flox mice were bred with EIIa-Cre transgenic mice (The Jackson Laboratory). Deletion of the KLK8 gene was confirmed by PCR analysis of genomic DNA using the following primers: (5’-GGACGTTGGAGTCACAGC-3’) and (5’-CCCAGGAGCAGAAGAGTG-3’). KLK8flox/flox; EIIa-Cre(+) mice (KLK8^-/-^) and age-matched KLK8flox/flox; EIIa-Cre(-) littermates (KLK8^+/+^) were used to investigate the effects of KLK8 deficiency. Mice with the same genotype were randomly assigned to a 1:1 allocation ratio for the control and endotoxemia groups. Endotoxemia was induced by intraperitoneal (i.p.) injection of purified LPS extracted from the membrane of Escherichia coli 0111:B4 (Sigma-Aldrich, Saint Louis, USA) at a dose of 30 mg/kg as described previously (Zhao et al., 2022).

For the construction of the CLP-induced sepsis model, anesthetized mice underwent a 1 cm midline incision of the abdomen. The cecum was exposed, ligated at a 0.5 cm point from the tip, and punctured with a 21-gauge needle (Zhang et al., 2021). After extruding a small quantity of fecal content, the cecum was returned to the abdominal cavity, and simple sutures were used to close the abdominal musculature and skin. Animals were rewarmed until fully conscious and provided free access to food and water. The sham group underwent the same procedures except for the CLP.

In an additional experiment, mice were randomly divided into two groups and treated with anti-KLK8 neutralizing antibodies (M081-3M2, MBL International; 20 μg/kg) or IgG isotype (MBL International; 20 μg/kg) via the tail vein. Thirty minutes after administration of anti-KLK8 neutralizing antibodies or IgG isotype, mice were injected intraperitoneally with LPS/saline or subjected to CLP/sham operation. The chosen dose of anti-KLK8 neutralizing antibodies was determined based on our preliminary experiments.

### 4.2 Isolation and culture of mouse lung vascular endothelial cells (MLVECs)

MLVECs were isolated from male C57BL/6 mice (3-4 weeks) as previously described (Zhao et al., 2022). Briefly, after anesthesia, the right ventricle of the mice was injected with PBS to clear blood from the lungs. Peripheral, subpleural lung tissues were cut into pieces and cultured in DMEM containing 20% fetal calf serum, 25 mM Hepes, 3.7 g/L NaHCO3, 5 mg/ml heparin, 1 mg/ml hydrocortisone, 80 mg/ml endothelial cell growth supplement from bovine brain, 5 mg/ml amphotericin and 0.01% ampicillin/streptomycin at 37°C with 5% CO2 for 60 hours. Subsequently, the diced tissue was removed, and the adherent cells were cultured in a basal culture medium. The MLVECs passaged between 3 and 4 times were used in experiments.

### 4.3 Immunofluorescence analysis

Paraffin sections (4 μm) of lung tissues were rehydrated and microwaved in citric acid buffer to retrieve antigens. After incubation with 10% BSA for 1 h, the sections were incubated with primary antibodies included: anti-CD31 (R&D Systems, AF3628), anti-KLK8 (Abcam, 232839), anti-Ki67 (Servicebio, GB111141) at a dilution of 1:100 at 4℃ overnight. After washes, sections were incubated with secondary antibodies conjugated with Alexa Fluor® 555 or Alexa Fluor® 488 at 37℃ for 1 h in the dark. Finally, nuclei were counterstained with 4’6-diamidino-2-phenylindole (DAPI) (Sigma-Aldrich). The fluorescent images were captured by Pannoramic MIDI (3D HISTECH, Budapest, Hungary) and analyzed using Image J software. The investigator performing immunofluorescence analysis was blinded to group allocation. For quantification, five high-power fields were analyzed in lung tissue sections taken from each mouse. We then determined the percentages of KLK8^+^/CD31^+^ cells, or Ki67^+^/CD31^+^ cells in total CD31^+^ cells.

### 4.4 Infection of adenovirus and lentivirus

Adenovirus (Ad) expressing KLK8 (Ad-KLK8) and lentivirus (Len) expressing FOXM1 (Len-FOXM1) were provided by Genechem (Shanghai, China). For intra-pulmonary adenovirus transfection, mice were intratracheally instilled with 1 × 10^8^ plaque-forming units (pfu) of Ad-KLK8 or Ad-Vector in 30 μl enhanced infection solution. The dose of adenovirus was used according to previous reports and our preliminary experiments(Mata-Espinosa et al., 2019).

### 4.5 3-[4,5-Dimethylthiazol-2-yl]-2,5-diphenyl tetrazolium bromide (MTT) assay

Cell viability was determined by MTT assay (Beyotime, China) based on the reduction of MTT by functional mitochondria to formazan(Du et al., 2021).

### 4.6 Fluorescein isothiocyanate-carboxymethyl (FITC)-dextran flux assay

The diffusion of FITC-dextran (10-kDa, Sigma-Aldrich) through the endothelial monolayer was determined to analyze the endothelial permeability in vitro as previously described (Zhao et al., 2022). MLVECs were seeded into the Transwell inserts (0.4 μm, 12 mm in diameter, Corning, NY, USA) and grown to confluence. After treatment, medium containing 1 mg/ml FITC-dextran was added to the top chamber of the Transwell for 1 h. Endothelial permeability was determined by measuring the amount of FITC-dextran in the lower compartment using a Synergy fluorescence plate reader (Bio-Tek, Cytation3, USA; excitation 485±20 nm, emission 528±20 nm).

### 4.7 Measurement of pulmonary transvascular permeability

The pulmonary transvascular albumin permeability was performed as previously described(Zhao et al., 2022). Briefly, evans blue–albumin (EBA) (40 mg/ml BSA with 1% evans blue dye, Sigma-Aldrich) was injected into the right jugular vein of the mice and allowed to circulate in the blood vessels for 45 min. Then intravascular evans blue was washed by PBS perfusion from the right ventricle for 2 min. Mouse lungs were excised, homogenized in 1 ml PBS, and extracted in 2 ml formamide (Sigma-Aldrich) overnight at 60 °C. Evans blue content was determined by OD620 using a spectrophotometer (Bio-Tek) of the formamide extract and normalized by body weight.

### 4.8 Lung histopathological evaluation

The pathological changes of lung tissues were observed under a light microscope after hematoxylin and eosin (H&E) staining. Lung injury score was assessed by two pathologists with expertise in lung pathology. The criteria for scoring lung injury were set up as previously described(Zhao et al., 2022) : 0 = normal tissue; 1 = tiny inflammatory change; 2 = mild to moderate inflammatory changes without marked damage in the lung architecture; 3 = moderate inflammatory injury with thickening of the alveolar septa; 4 = moderate to severe inflammatory injury with the formation of nodules or areas of pneumonitis; and 5 = severe inflammatory injury with total obliteration of the field. The mean score was reported per section.

### 4.9 Open Field Test (OFT)

An OFT was performed to evaluate LPS-induced behavioral deficits in mice(Zhang et al., 2018, Li et al., 2020). Briefly, mice were placed at the center of the open field (40 cm length × 40 cm width × 30 cm height) and allowed to freely explore for five minutes. A video-computerized tracking system (ANY-maze, Stoelting CO, EUA) was used to track and record the movement of mice. The parameters evaluated were the total, central and peripheral distances traveled, the duration of time spent in the central and peripheral areas, and the frequency of crossing squares. The chamber was cleaned with 75% ethanol before the next test.

### 4.10 Transfection of small interfering RNA (siRNA)

The siRNAs for KLK8 were provided by GenePharma Corporation (Shanghai, China). The sequences for mouse KLK8 siRNA are: 5’-CCUGGAUCAAGAAGACCAUTT-3’ and 5’-AUGGUCUUCUUGAUCCAGGTT-3’. Negative control siRNA was scrambled sequence without any specific target: 5’-UUCUCCGAACGUGUCACGUTT-3’ and 5’-ACGUGACACGUUCGGAGAATT-3’. Transfection of siRNA in MLVECs was performed by using the Lipofectamine TM3000 (Thermo Fisher) according to the manufacturer’s instructions.

### 4.11 5-ethynyl-2’-deoxyuridine (EdU) cell proliferation assay

MLVECs proliferation was determined using BeyoClick™ EdU-594 Kit (Beyotime). Briefly, MLVECs (1 × 10^5^/well) were seeded into 24-well plates. After treatment, the cells were incubated at 37 °C and 5% CO2 in medium containing 10 μM EdU for 2 h, and then fixed with 4% paraformaldehyde for 15 min. After removing the supernatant, cells were treated with PBS with 0.3% Triton X-100 and then stained with the Click Additive Solution at 37°C for 30 min in the dark. The nucleus was stained with DAPI. Fluorescent images were taken by a Zeiss LSM 710 laser confocal microscope (Zeiss). Cell proliferation was determined as the ratio of the number of EdU-positive nuclei to that of DAPI-positive nuclei.

### 4.12 Western blot analysis

Proteins were extracted from lung tissues or MLVECs using RIPA (Beyotime, China) containing protease and phosphatase inhibitor cocktail (Beyotime) according to the manufacturer’s instructions. Protein concentrations were determined by BCA protein assay kit (Beyotime). Equal amounts of proteins (30 μg) were separated with 10% SDS-PAGE, transferred to PVDF membrane (Millipore, Billerica, MA, USA), and then blocked with 5% non-fat milk. Membranes were incubated overnight with specific primary antibodies against KLK8 (Abcam Cat# ab232839), CD31 (R&D Systems Cat# AF3628), ZO-1 (Proteintech Cat# 21773-1-AP), intercellular adhesion molecule 1 (ICAM-1) (Proteintech Cat# 10020-1-AP), FOXM1 (Santa Cruz, Cat# sc-376471), phospho-Akt (Ser473, Proteintech Cat# 66444-1-Ig), Akt (Proteintech Cat# 10176-2-AP), phospho-mTOR (Ser2448, CST, Cat# 2971), mTOR (CST, Cat# 2983), phospho-p70 S6K (Thr389, Proteintech Cat# 28735-1-AP), p70 S6K (Proteintech Cat# 14485-1-AP), vascular endothelial (VE)-cadherin (Santa Cruz, Cat# sc-52751), and β-actin (Sigma-Aldrich Cat# A5441). The antibody-reactive bands were visualized using an enhanced chemiluminescence Western blotting detection system (Millipore).

### 4.13 Immunoprecipitation

MLVECs or lung tissues were lysed with cold RIPA lysis buffer (Beyotime) containing 1% Proteinase Inhibitor Cocktail (Sigma-Aldrich) and proteasome inhibitor (N-ethylmaleimide, at 20mM). Cell lysates were incubated on ice for 2 hours, and cell debris was removed by centrifugation. The clarified supernatants were incubated with antibodies against KLK8 or VE-cadherin at 4 ℃ for 16 hours with gentle rotation. IgG was used as control for nonspecific interaction. Protein A and G sepharose beads (Beyotime) were added and incubated at 4℃ for an additional 3 hours, and the immune complexes were washed four times. The final precipitate was boiled in a protein loading buffer for 5 minutes and eluted on 10% SDS-PAGE for Western blot analysis using the respective antibodies.

### 4.14 Quantitative Real-time RT-PCR

For RNA isolation and reverse transcription (RT) in lung tissue and MLVECs, Total RNA was isolated using Trizol reagent (Vazyme), and then performed with 4 μg RNA of total RNA for RT, Quantitative PCR was operated using SYBR® Premix Ex Taq (Vazyme) and operated with a StepOne Plus (Applied Biosystem, Foster City, CA) apparatus. Endogenous Ct values of β-actin were used as a control. The primer sequences used in this study were provided in Supplemental Table 1.

### 4.15 RNA Sequencing (RNA-seq) and bioinformatic analysis

MLVECs for RNA-seq experiments were seeded into 6-well plates, and infected with Ad-KLK8 or Ad-Vector at a multiplicity of infection (MOI) of 3 in serum-free medium for 48 h. After treatment, MLVECs and the culture medium were collected for RNA-seq analysis or measurement of KLK8 protein levels, respectively. As shown in Fig. 6E, it was found that infection of Ad-KLK8 significantly increased KLK8 release in the culture medium, indicating the successful overexpression of KLK8 in MLVECs. Total RNA of MLVECs was extracted using Trizol reagent (Vazyme). RNA-seq analysis was performed by OE Biotech (Shanghai, China). The constructed library was qualified with the Agilent 2100 Bioanalyzer, and the Illumina HiSeq™ 2500 platform was used for sequencing. The gene expression was measured in units of fragments per kilobase of exon model per million mapped reads (FPKM). A p-value with correction for multiple testing using the Benjamini-Hochberg (BH) procedure, BH-corrected false-discovery rate (FDR) < 0.05, and FoldChange (FC) ≥2 or FC ≤ 0.5 (|LogFC |≥ 1) were set as the threshold for significant differential expression. Further, gene set enrichment analysis (GSEA, version 4.3.0) was used to analyze pathway enrichment (http://software.broadinstitute.org/gsea/). Upstream regulator analysis was performed using Ingenuity Pathway Analysis (IPA, QIAGEN, Redwood City, CA). All the RNA-Seq data were deposited in the National Center for Biotechnology Information (NCBI) GEO database under the accession number GSE216969.

### 4.16 Statistical Analysis

Statistical analyses were done with SPSS 26.0 (SPSS Inc., Chicago, USA). The results are expressed as mean ± SEM. Normal distribution was assessed by the Shapiro-Wilk test. Statistical significance in experiments comparing only two groups was determined by a two-tailed Student’s t-test. The comparison among multiple groups was estimated by one-way analysis of variance followed by post hoc analysis using the Student-Newman-Keuls test. Kaplan–Meier test was used to determine statistical significance between survival curves. A P value of < 0.05 was considered significant.

## Availability of data and materials

Data and materials are available upon reasonable request to the corresponding author Professor Zhu.

## Abbreviations

KLKs: kallikrein-related peptidases
KLK1: kallikrein-related peptidase 1
KLK8: kallikrein-related peptidase 8
LPS: lipopolysaccharide
Ad-KLK8: Adenovirus KLK8
VE-Cadherin: vascular endothelial cadherin
MODS: multiple organ dysfunction syndrome
MLVECs: mouse lung vascular endothelial cells
EndMT: endothelial-to-mesenchymal transition
CLP: cecal ligation and puncture
Len: lentivirus
Ad: Adenovirus
pfu: plaque-forming units
FITC: Fluorescein isothiocyanate-carboxymethyl
MTT: 3-[4,5-Dimethylthiazol-2-yl]-2,5-diphenyl tetrazolium bromide
EBA: evans blue–albumin
H&E: hematoxylin and eosin
OFT: Open Field Test
siRNA: small interfering RNA
EdU: 5-ethynyl-2’-deoxyuridine
RT: reverse transcription
RNA-seq: RNA Sequencing
IPA: ingenuity pathway analysis
DEGs: differentially expressed genes
MOI: multiplicity of infection
GO: Gene Ontology
KEGG: Kyoto Encyclopedia of Genes and Genomes
ICAM-1: intercellular adhesion molecule 1
mTOR: Mammalian target of rapamycin
MTORC1: The mechanistic target of rapamycin complex 1
FOXM1: Forkhead box M1

## Acknowledgements

We thank Dr. Jian-Kui Du (Navy Medical University) for providing the lentivirus expressing VE-Cadherin.

## Competing interests

The authors declare no competing interests.

## Ethical approval and consent to participate

All animal experiments were performed in compliance with the guidelines provided by the Institutional Animal Care and Use Committee’s guidelines of Navy Medical University, Shanghai, China. The Committee on the Ethics of Animal Experiments at Navy Medical University approved this study.

## Consent for publication

Not applicable.

## Funding

This research was supported by National Natural Science Foundation of China (No. 32171124, 31871156, 31971101, 32271180, 82272229, 81471852) and Hunan Provincial Natural Science Foundation (2021JJ31058).

## Author contributions

X.Z. and Y.L. conceived this project. Y.Z., H.J. and F.H. performed the major experiments. Q.X., H.Z. and D.L. contributed to animal operations and lung tissue collection. J.W. and D.X. contributed to the western blot and qPCR analysis and corresponding analysis of data. X.Z., Y.L., L.J. and P.X. contributed to the overseen and supervision of the whole project. X.Z. and Y.L. provided the fund for this study. X.Z. and Y.Z. wrote and revised the manuscript. All authors read and approved the final manuscript.

